# Pathogenic CD8 T cell responses are driven by neutrophil-mediated hypoxia in cutaneous leishmaniasis

**DOI:** 10.1101/2023.10.18.562926

**Authors:** Erin A. Fowler, Camila Farias Amorim, Klauss Mostacada, Allison Yan, Laís Amorim Sacramento, Rae A. Stanco, Emily D. S. Hales, Aditi Varkey, Wenjing Zong, Gary D. Wu, Camila I. de Oliveira, Patrick L. Collins, Fernanda O. Novais

**Author notes:** Corresponding author: Fernanda O. Novais, The Ohio State University, College of Medicine 720 BRT 460 W 12th Ave, Columbus OH 43210. Division of Gastroenterology/Hepatology/Nutrition, Children’s Hospital of Philadelphia; Philadelphia, USA. Conflict of Interest: The authors have declared no conflict of interest exists.

## Abstract

Cutaneous leishmaniasis caused by *Leishmania* parasites exhibits a wide range of clinical manifestations. Although parasites influence disease severity, cytolytic CD8 T cell responses mediate disease. While these responses originate in the lymph node, we find that expression of the cytolytic effector molecule granzyme B is restricted to lesional CD8 T cells in *Leishmania*- infected mice, suggesting that local cues within inflamed skin induce cytolytic function. Expression of Blimp-1 (*Prdm1*), a transcription factor necessary for cytolytic CD8 T cell differentiation, is driven by hypoxia within the inflamed skin. Hypoxia is further enhanced by the recruitment of neutrophils that consume oxygen to produce reactive oxygen species, ultimately increasing granzyme B expression in CD8 T cells. Importantly, lesions from cutaneous leishmaniasis patients exhibit hypoxia transcription signatures that correlate with the presence of neutrophils. Thus, targeting hypoxia-driven signals that support local differentiation of cytolytic CD8 T cells may improve the prognosis for patients with cutaneous leishmaniasis, as well as other inflammatory skin diseases where cytolytic CD8 T cells contribute to pathogenesis.

## INTRODUCTION

Cutaneous leishmaniasis is caused by *Leishmania* parasites and exhibits a wide range of clinical manifestations, from self-healing lesions to chronic debilitating infections (*1*). No vaccines exist for leishmaniasis, and the anti-parasitic drugs are often ineffective (*2*). Although the parasites influence disease severity, immune responses are often the major driver of disease. For example, CD8 T cells in lesions are recruited to target parasite-infected cells and kill infected host cells through granule-mediated cytotoxicity. However, the destruction of infected cells does not kill *Leishmania*, but leads to the release of parasites that metastasize to distant cutaneous sites (*3*). In addition, cell death induced by CD8 T cells activates the NLRP3 inflammasome, IL- 1β release, chronic neutrophil recruitment, and enhanced inflammation (*3–6*). The importance of this pathway is supported by our previous analysis of lesions from patients, in which granzyme B (*GZMB*), perforin (*PRF1*), and *IL1B* expression were associated with treatment failure (*7*). While both murine and human studies underscore the pathogenic consequences of cytotoxicity, the microenvironmental cues and downstream mechanisms that drive cytolytic programs in lesional CD8 T cells remain unknown.

In response to *Leishmania* infection, CD8 T cells are activated and expand in draining lymph nodes (dLNs). The activated cells then travel to inflammatory sites, where they kill infected cells. While infections also induce differentiation of effector CD4 T cells in secondary lymphoid organs, CD8 T cells with cytotoxic effector function are generally absent in LNs (8–11). A prior study implicated PD-1 ligation on dLN CD8 T cells in suppressing their expression of granzyme B, a process that is likely advantageous since it would limit the killing of antigen-presenting cells (11). While such findings suggest that CD8 T cell cytolytic effector function is only triggered once CD8 T cells enter inflammatory sites (12, 13), the local cues that drive this response remain unclear. As the CD8 T cell cytolytic program can have both positive and negative consequences for the host (14–33), understanding the mechanisms that drive cytotoxic CD8 T cell programs remains an important goal for unraveling pathogenesis of numerous diseases such as alopecia, vitiligo, bullous pemphigoid, Steven-Johnson syndrome and toxic epidermal necrolysis, where cytolytic CD8 T cells contribute to pathogenesis (15, 32, 34–37).

One potential mechanism relevant to inflamed tissues is hypoxia, which occurs when oxygen consumption by infiltrating cells exceeds the supply (38) and differentially impacts immune cell types. In *Leishmania* infection, hypoxia has been studied exclusively in macrophages and dendritic cells (39, 40), where it can either promote or block effector responses, depending on the parasite species and the cell type (41–49). Here, we find that hypoxia within *Leishmania*-infected lesions directly promotes cytolytic effector function in newly recruited CD8 T cells. Mechanistically, we identify a feed-forward loop, initiating with early recruitment of neutrophils that consume oxygen to produce reactive oxygen species (ROS), which promotes a persistently hypoxic environment, elevates the number of cytolytic CD8 T cells, perpetuates chronic neutrophil recruitment, and exacerbates tissue damage. Thus, our findings indicate that targeting cytolytic CD8 T cells driven by hypoxic microenvironments will improve the prognosis for patients with cutaneous leishmaniasis and other inflammatory skin diseases.

## RESULTS

### CD8 T cells express granzyme B in leishmanial lesions but not in the draining lymph nodes

Cytotoxic CD8 T cells are pathogenic in cutaneous leishmaniasis (3, 4, 6, 7, 50–54), but their anatomical distribution and mechanisms that regulate their induction remain unclear. In this regard, parasites are found in both dLNs and skin lesions. To determine if CD8 T cell cytotoxic effector function is present in both tissues, we assessed the expression of the cytotoxic marker granzyme B. C57BL/6 mice were infected with *Leishmania* in the ear, and two weeks post- infection, the frequency of granzyme B^pos^ antigen-experienced (CD44^high^) CD8 T cells was assessed by flow cytometry. While granzyme B-expressing CD8 T cells were abundant in the infected skin, granzyme B expression was nearly absent in dLNs (Fig 1a). To determine if granzyme B^pos^ CD8 T cells were preferentially recruited to the skin, we treated mice with FTY720, a sphingosine 1-phosphate receptor agonist that blocks the egress of T cells from the dLNs. If granzyme B^pos^ CD8 T cells are preferentially recruited to cutaneous lesions, granzyme B^pos^ CD8 T cells should accumulate in the dLNs of FTY720-treated mice. However, there was no difference in the frequency of granzyme B^pos^ CD8 T cells in the dLNs of mice treated with FTY720 or vehicle control (Fig. 1b), suggesting that rather than preferential recruitment, exposure to the lesion microenvironment induces a cytotoxic profile in CD8 T cells. To directly test this, we infected C57BL/6 CD45.1 or CD45.2 congenic mice with *Leishmania*, and purified CD8 T cells from dLNs of CD45.2 donor mice (granzyme B^neg^) three weeks post-infection. Donor dLN CD8 T cells were then transferred directly into the lesions of recipient CD45.1 infected animals (Fig. 1c - schematic). Notably, we found that granzyme B^neg^ CD8 T cells became granzyme B-expressing CD8 T cells in the lesions (Fig. 1c). In contrast, CD8 T cells that migrated back to dLNs remained granzyme B^neg^ (Fig. 1c). These results demonstrated that the lesion microenvironment triggers cytotoxic programs in CD8 T cells.

**Fig 1.**
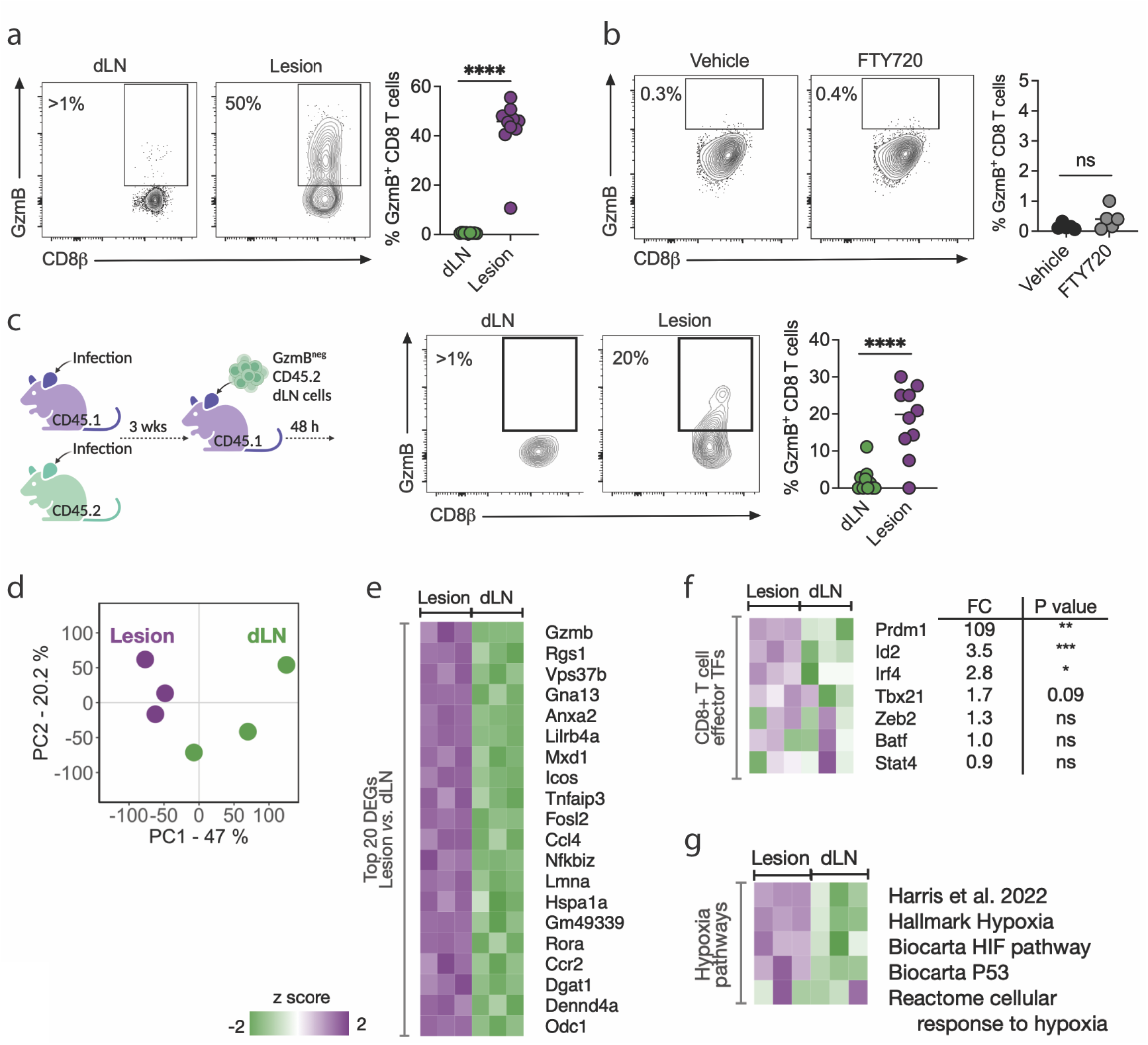
CD8 T cell function is tissue-specific. (a) Granzyme B (GzmB) expression in CD8 T cells from dLNs and lesions from C57BL/6 mice in week two post-infection with *L. major*. Representative data from more than three experiments with at least four mice per experiment. (b) Granzyme B expression in dLN CD8 T cells from *L. major* infected C57BL/6 mice treated with FTY720 or vehicle daily 12 days post-infection for 10 days. Data from one independent experiment with five mice per group. (c - left) Schematic representation of the transfer of CD45.2 dLN purified CD8 T cells into the lesions of CD45.1 recipients, both infected for three weeks. (c - right) Granzyme B expression in donor CD8 T cells from dLNs and lesions from recipient mice 48 hours post-transfer. Data from 2 experiments independent experiments combined. (d-g) RNA-seq analyses from antigen-experienced (CD44^high^) CD8 T cells purified from dLNs and lesions of *L. major*-infected mice in week five. (d) Principal component analysis showing principal component 1 (PC1) and PC2. (e) Heat map of top 20 differentially expressed genes (DEGs) between purified CD8 T cells from the lesions compared to dLN (FDR>0.05 and FC=1.5). (f) Expression of genes encoding for transcription factors associated with effector-like T cell functions is represented in a heat map, as well as fold-change (FC) between lesional and dLN CD8 T cells. (g) Heatmap of hypoxia-related pathways enriched in lesional and dLN CD8 T cells. *P ≤ 0.05, **P ≤ 0.01, ***P ≤ 0.001 and ****P ≤ 0.0001. ns = non-significant. FDR = false discovery rates. FC = fold change.

To determine the mechanisms by which CD8 T cell cytotoxic programs are induced in lesions, we performed RNA-seq analysis on purified antigen-experienced CD8 T cells from dLNs and lesions of mice infected with *Leishmania* for five weeks. Principal component analysis (PCA) showed that almost half (Principal component 1, PC1 47%) of the differences on the whole transcriptional profiles were associated with the organ from which the CD8 T cells were derived (Fig. 1d). Differentially expressed gene (DEG) analysis with thresholds of false discovery rate (FDR) < 0.05 and fold change > 1.5 between tissues revealed 118 genes overexpressed in CD8 T cells from lesions compared to dLNs and 26 DEGs overexpressed in dLNs compared to lesions (Supp. Table 1 and Suppl. Fig. 1). *Gzmb* was the top DEG in CD8 T cells from lesions compared to dLNs, FDR=0.002 (Fig. 1e). Several transcription factors play important roles in effector CD8 T cell biology, including BATF, ID-2, IRF4, Stat4, T-bet, Zeb2, and Blimp-1 (encoded by *Prdm1*) (55–57), and we found that *Id2*, *Irf4*, and *Prdm1* expressions were significantly higher in lesions than in dLNs, and *Prdm1* had the most significant fold change between tissues (>100 higher in lesions) (Fig. 1f). To further investigate the signals received by CD8 T cells within lesions, we performed Gene Set Enrichment Analysis (GSEA), which exhibited hypoxia-related signatures, as assessed by several pathway databases (Hallmark, Biocarta, and Reactome), as well as the Harris et al. hypoxia-specific pathway described in (58), (Fig. 1g and Supp. Table 2). These data indicate that, after their exit from dLNs, CD8 T cells recruited to cutaneous leishmaniasis lesions are exposed to hypoxic conditions, inducing the expression of key cytotoxicity effector mediators, including *Prdm1*.

### Cutaneous leishmaniasis lesions are hypoxic and alter CD8 T cell function

Normal skin is naturally low in oxygen (59, 60), and inflammatory environments are frequently hypoxic. To evaluate hypoxia in *Leishmania*-infected lesions, we employed pimonidazole, a 2-nitroimidazole reporter molecule that is reductively activated and forms covalent bonds with macromolecules, specifically in hypoxic cells. Mice were injected with pimonidazole two weeks after *Leishmania* infection and one hour before euthanasia.

Pimonidazole staining was assessed in infected and contralateral ears by confocal microscopy. As expected, we found pimonidazole staining within the epidermis, dermis, and hair follicles of naïve mice (Fig. 2a and 2b – top images). Within lesions, we observed enlargement of the epidermis and dermis associated with robust pimonidazole staining (Fig. 2a and 2b – bottom images). Surprisingly, pimonidazole staining was absent from ulcerated areas. Representative images in lower (Fig. 2a and Supp. Fig. 2a) and higher (Fig. 2b and Supp. Fig. 2b) magnification depict the differences in pimonidazole staining distribution between the naïve and infected tissue.

**Fig 2.**
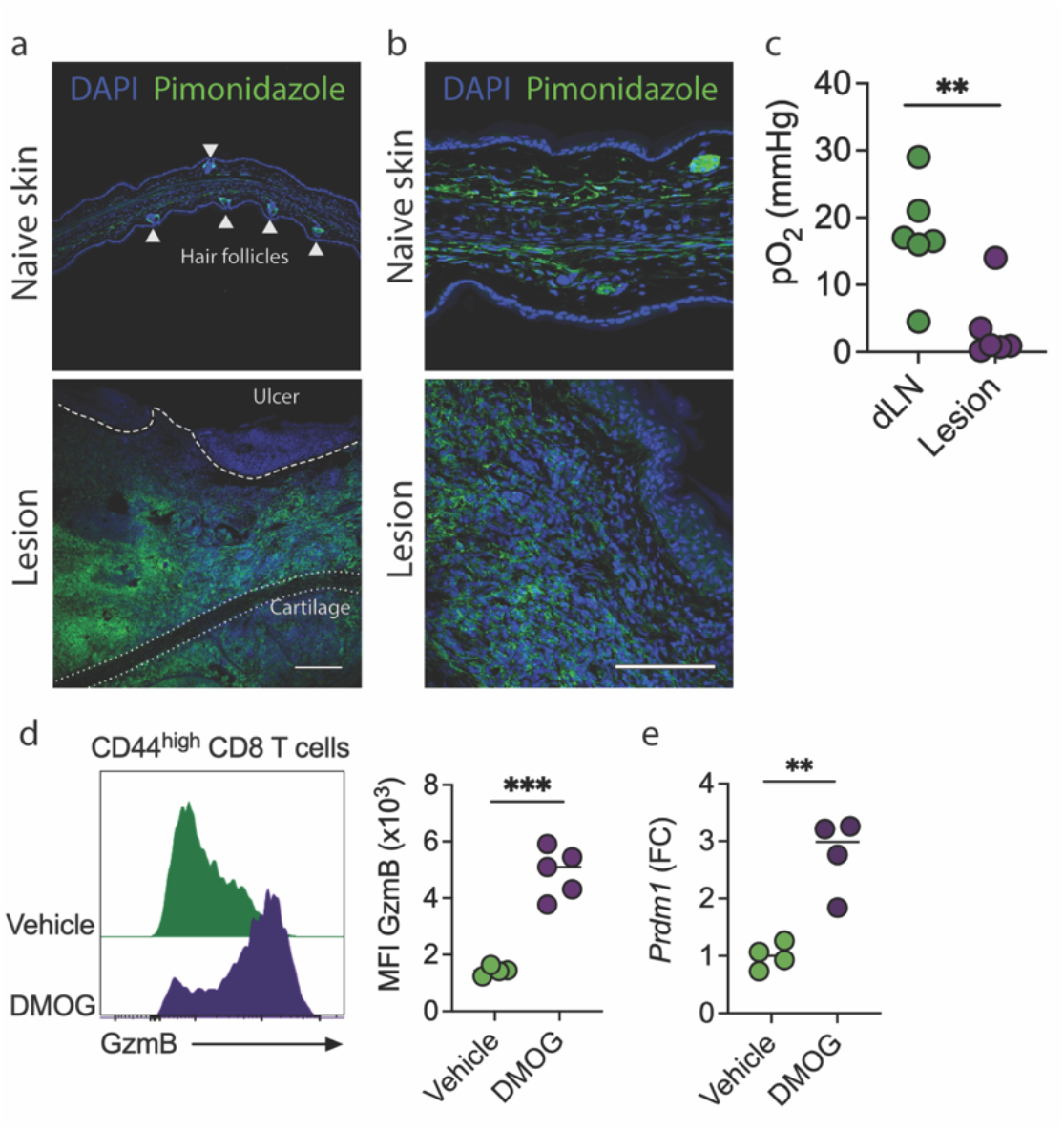
Hypoxia induces granzyme B and *Prdm1* expression in CD8 T cells. (a-b) Immunofluorescence staining and confocal microscopy of horizontal sections from lesions of *L. major* C57BL/6-infected mice for two weeks. Mice received pimonidazole one hour before euthanasia. The top panels show contralateral ears (naïve skin), and the bottom panels are from infected ears (lesions). Pimonidazole (green) and nuclear stain DAPI (blue). Representative image from two naïve and three infected ears. Bars (a) 200 μm and (b) 100 μm. (c) Partial pressure of oxygen in mmHg in dLNs and lesions of C57BL/6 mice infected with *L. major* for two weeks. Data from two independent experiments combined. (d) Granzyme B (GzmB) expression in CD8 T cells from dLNs of C57BL/6 mice infected with *L. major* cultured with Dimethyloxalylglycine (DMOG) or vehicle. (e) Fold-change (FC) of *Prdm1* mRNA over the average expression of vehicle-treated cells measured by qPCR in DMOG or vehicle-treated purified CD8 T cells. **P ≤ 0.01 and ***P ≤ 0.001

To compare O2 levels in lesions and dLNs, we infected mice with *Leishmania,* and two weeks post-infection, mice received Oxyphor G4, a phosphorescent probe quenched by oxygen (61). Infected skin exhibited a significant decrease in oxygen compared to dLNs (Fig. 2c). These data directly demonstrated skin lesions present a hypoxic microenvironment to CD8 T cells upon their recruitment from dLNs.

We tested whether exposure to hypoxia was sufficient to promote granzyme B expression in dLN CD8 T cells. We isolated dLN cells from infected mice and stimulated them with anti- CD3 and anti-CD28 in the presence of Dimethyloxalylglycine (DMOG), a compound that mimics hypoxia at normal oxygen tension. As shown in Fig. 2d, DMOG induced the expression of granzyme B in CD8 T cells, further demonstrating that hypoxia promotes upregulation of their cytolytic program. Notably, DMOG treatment also increased *Prdm1* mRNA expression compared to vehicle-treated CD8 T cells (Fig. 2e). Collectively, these data demonstrate that the hypoxic state of lesions stimulates differentiation of recruited CD8 T cells into cytotoxic effectors.

### Blimp-1 expression is restricted to granzyme B expressing CD8 T cells

Blimp-1 deficient CD8 T cells produce less granzyme B (62), so we next sought to test cause-effect relationships between hypoxia, Blimp-1, and cytotoxicity. To address this question, we infected Blimp-1 Yellow Fluorescent Protein (YFP) reporter mice with *Leishmania* and assessed Blimp-1 expression in CD8 T cells. We found that Blimp-1 was highly expressed in lesional CD8 T cells but not in CD8 T cells from the dLNs (Fig. 3a). We also found that Blimp-1 expression was significantly higher in CD8 T cells that express granzyme B (Fig. 3b), suggesting a link between Blimp-1 and cytotoxicity. Finally, to assess if granzyme B expression in lesions resulted from hypoxic induction of Blimp-1, we used mice expressing a fusion protein bearing an oxygen-dependent degradation (ODD) domain from human HIF1A (hypoxia-inducible factor 1, alpha subunit) fused to a tamoxifen-inducible Cre recombinase gene (cre/ERT2) (63). When expressed under normoxic conditions, the O2CreER fusion protein is degraded rapidly but is stabilized by hypoxia when combined with tamoxifen injection. These mice were crossed with Blimp-1^flox/flox^ mice (here called ODD^cre^Blimp-1^flox/flox^ mice) to generate mice in which Blimp-1 expression is deleted specifically in hypoxic cells upon tamoxifen injection (Fig. 3c). Since the skin has significantly less oxygen than dLNs, these mice provide lesion-specific deletion of Blimp-1, thus ensuring the preservation of Blimp-1-dependent development of effector T cells in dLNs (normoxic) and their recruitment to the inflamed tissue. *Leishmania*-infected ODD^cre^Blimp-1^flox/flox^ mice were treated with tamoxifen or left untreated for one week, and granzyme B expression was evaluated. We found that specific deletion of Blimp-1 in hypoxic cells significantly decreased granzyme B expression in lesional CD8 T cells compared to untreated mice (Fig. 3d). Taken together, these results directly demonstrate that upregulation of Blimp-1 by the hypoxic environment leads to increased granzyme B expression and CD8 T cell cytotoxicity in leishmanial lesions.

**Fig 3.**
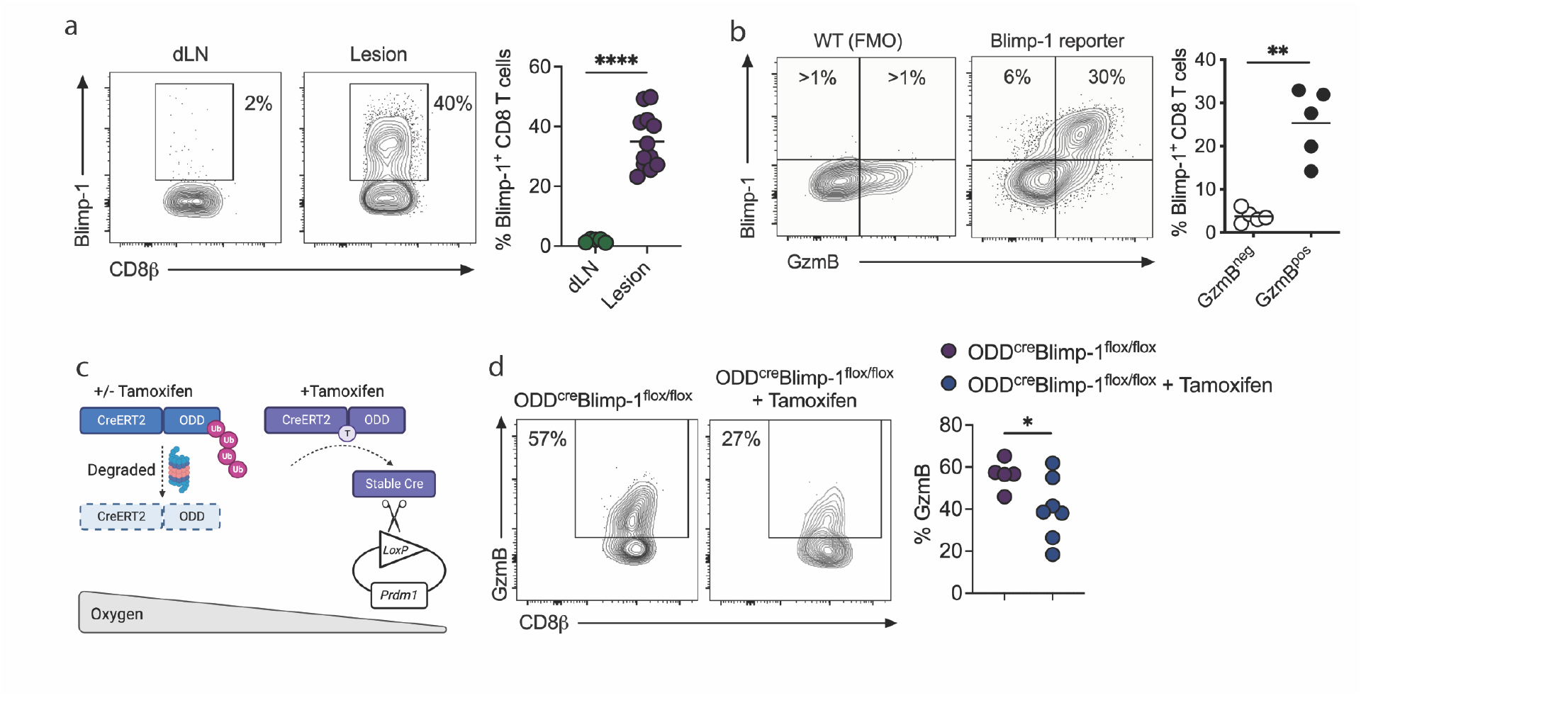
Blimp-1 expression is induced by hypoxia within skin lesions. (a) Blimp-1 YFP reporter mice expression in CD8 T cells from dLNs and lesions from Blimp-1 reporter mice infected with *L. major* for one week. Data from two independent experiments combined. (b) Granzyme B (GzmB) and Blimp-1 YFP expression in CD8 T cells from dLNs and lesions from Blimp-1 reporter mice infected with *L. major* for one week. (c) Schematic representation of OOD^cre^Blimp-1^flox/flox^ mice. (d) OOD^cre^Blimp-1^flox/flox^ mice were infected with *L. major* for one week when daily tamoxifen injections commenced. Mice were euthanized at week two post-infection. GzmB expression of lesional CD8 T cells in tamoxifen-treated and untreated mice. Data from two independent experiments combined. *P ≤ 0.05, **P ≤ 0.01, and ****P ≤ 0.0001.

### Blimp-1 expression is necessary for CD8 T cells to mediate disease

Based on our data, we predicted that Blimp-1 expression was required for cytotoxic CD8 T cell-mediated disease in leishmaniasis. Since Blimp-1 plays a vital role in CD4 T cell subsets (64) and regulates IL-10 production (65), total T cell deletion of Blimp-1 would complicate interpretations. Therefore, we used our well-characterized mouse model of chronic leishmaniasis in which severe disease develops following *Leishmania* infection of RAG-deficient mice reconstituted with CD8 T cells (3, 4, 66, 67). RAG-deficient mice infected with *Leishmania* were reconstituted with WT or CD8 T cells lacking Blimp-1 expression (here called Blimp-1^cKO^) or control animals receiving no T cells. As described previously (3), RAG-deficient mice reconstituted with WT CD8 T cells developed severe pathology characterized by granzyme B-expressing CD8 T cells in lesions, while RAG-deficient mice that received no cells showed no signs of pathology (Fig. 4a). Importantly, RAG-deficient mice reconstituted with Blimp-1^cKO^ CD8 T cells had minimal disease (Fig. 4a). As expected, there were similar numbers of parasites (Fig. 4b) and a similar frequency of CD8 T cells (Fig. 4c) in lesions of RAG-deficient mice reconstituted with WT and Blimp-1^cKO^ CD8 T cells. Notably, granzyme B expression was significantly diminished in cells lacking Blimp-1 (Fig. 4d). Collectively, these data indicate that Blimp-1 expression is driven by the hypoxic microenvironment of the lesion, triggering granzyme B expression and CD8 T cells-driven pathology.

**Fig 4.**
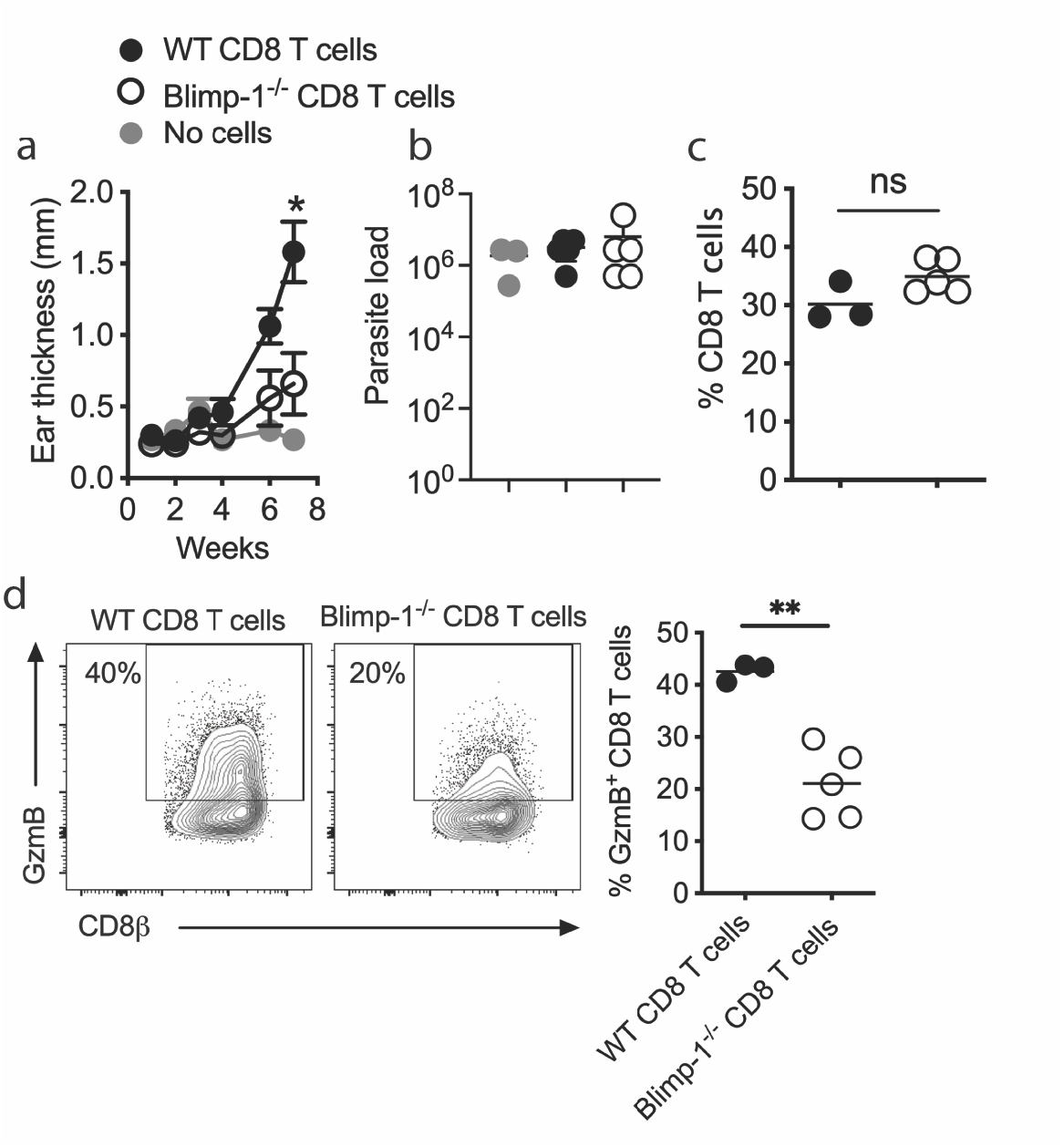
Blimp-1 expression is required for CD8 T cell-mediated disease. RAG^−/−^ mice were infected with *L. braziliensis* and reconstituted with CD8 T cells purified from WT or Blimp-1^cKO^ mice. (a) Ear thickness and (b) parasite numbers in lesions at seven weeks of infection. (c) CD8 T cell frequency and (d) GzmB expression by CD8 T cells in lesions were assessed directly ex vivo by flow cytometry seven weeks after infection. Data represent 3 individual experiments with 3-5 mice per group. *P ≤ 0.05 and **P ≤ 0.01. ns = non- significant

### Neutrophils generate the hypoxic microenvironment of cutaneous leishmaniasis lesions

Hypoxia occurs in tissues when oxygen supply does not meet demand, including scenarios characterized by defective tissue vascularization or increased demand by cells present within a tissue (38). *Leishmania*-infected lesions are highly vascularized (68), so we hypothesized that oxygen consumption by inflammatory cells recruited to cutaneous leishmaniasis lesions promoted hypoxia. Neutrophils are the first cells recruited to skin upon *Leishmania* infection (69), but their recruitment does not control *L. major* parasites. Indeed, their chronic presence is associated with worsening disease (3, 4, 70–74). Therefore, we tested whether neutrophil recruitment contributed to the hypoxic state of inflamed skin. *Leishmania*- infected mice were injected with pimonidazole 1 hour before euthanasia at multiple time points after infection: 1) soon after parasite challenge (2 and 5 hours), 2) when lesions started to develop, and CD8 T cells were present and expressed granzyme B (2 weeks), and 3) when lesions began to heal (9 weeks). As expected (69), neutrophils were quickly recruited to *Leishmania*-infected lesions and decreased in frequency when lesions resolved (Fig. 5a).

**Fig 5.**
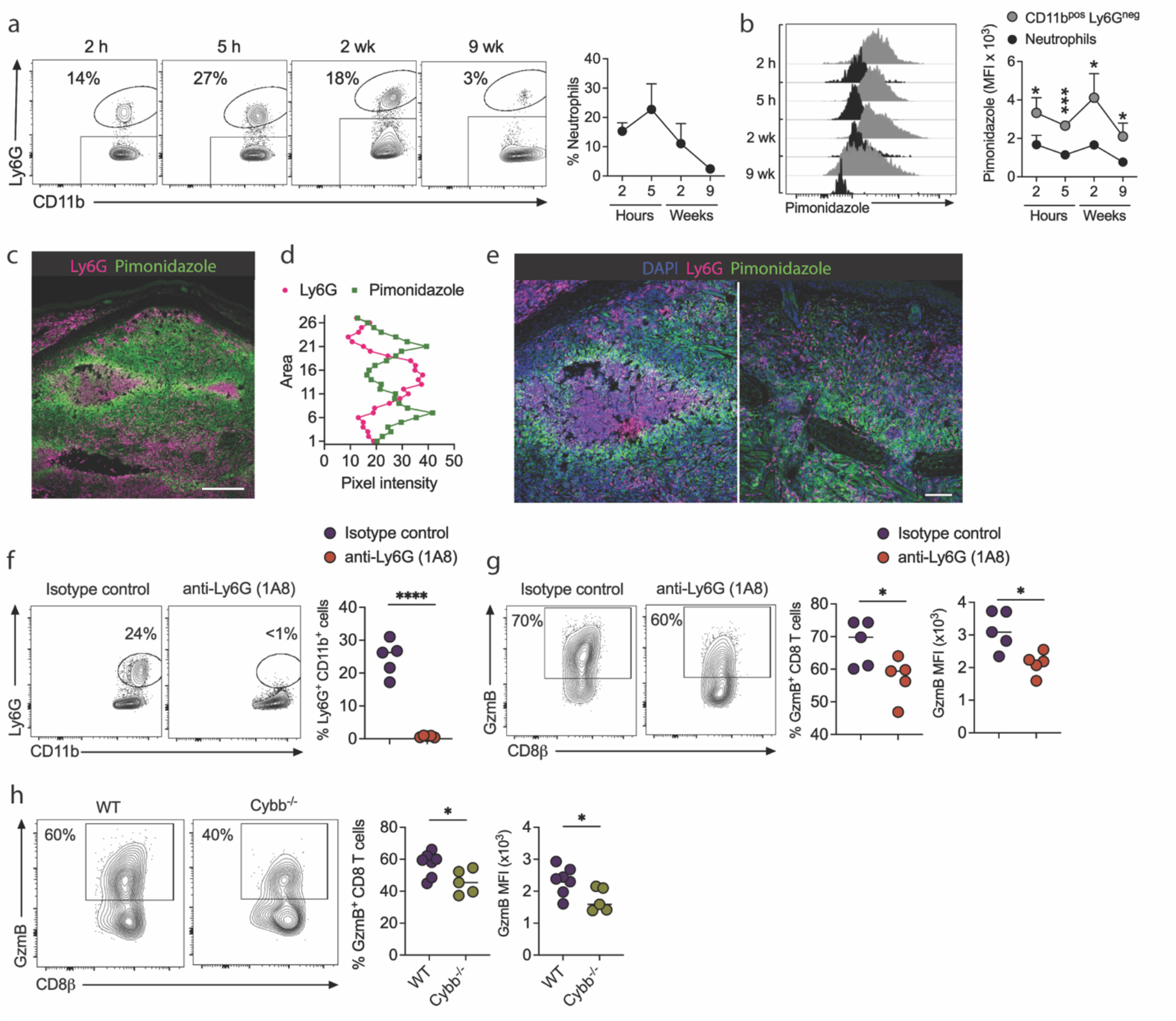
Neutrophils consume oxygen from lesions and stimulate granzyme B expression in CD8 T cells. (a-b) C57BL/6 mice infected with *L. major* received pimonidazole one hour before euthanasia at two hours, five hours, two weeks, and nine weeks. (a) Frequency of neutrophils in lesions and (b) pimonidazole expression in neutrophils (CD11b^+^Ly6G^pos^) and other myeloid cells (CD11b^+^Ly6G^neg^) in lesions. (c) Confocal microscopy of one representative image of Ly6G (pink) and pimonidazole (green) staining in the skin infected for two weeks. Scale bar = 200 μm. (d) Pixel intensity of pimonidazole and Ly6G based on 27 regions (Fig. S3). (e) Confocal microscopy of two representative images of nuclear (DAPI) staining (blue), Ly6G (pink), and pimonidazole (green) in lesions infected for two weeks. Scale bar = 100 μm. (f-g) C57BL/6 mice infected with *L. major* were injected every three days with anti-Ly6G (clone 1A8) or isotype control until euthanasia at two weeks post-infection. Frequency of (f) neutrophils in lesions. (g) Granzyme B (GzmB) frequency and mean fluorescence intensity (MFI) in CD8 T cells in lesions. (h) Granzyme B frequency and MFI in CD8 T cells in the lesions of WT or Cybb^-/-^ mice infected with *L. major* for two weeks. *P ≤ 0.05, ***P ≤ 0.001 and ****P ≤ 0.0001.

Surprisingly, neutrophil (CD11b^pos^Ly6G^pos^) staining by pimonidazole was significantly lower compared to other myeloid cells (CD11b^pos^Ly6G^neg^) in the skin at all time points analyzed (Fig. 5b). To confirm these observations, lesions from pimonidazole-injected mice were stained for Ly6G (clone 1A8), revealing intense neutrophil recruitment in ulcered regions in the epidermis and clusters of neutrophils within the skin dermis (Fig. 5c and Fig. S3a). Notably, regions without neutrophils had significantly more pimonidazole staining than those containing neutrophil clusters. We also observed that pimonidazole staining in regions surrounding neutrophil clusters was more intense. Quantitative assessment of pixel intensities (Supp. Fig. 3b) revealed a negative correlation between pimonidazole and Ly6G expression (Fig. 5d). Higher magnification of regions with intense neutrophil recruitment at two different time points showed a lack of pimonidazole staining in neutrophils (Fig. 5e and Supp. Fig. 3c).

These results suggest that areas surrounding neutrophils are relatively hypoxic. Accordingly, we hypothesized that neutrophils were responsible for promoting the hypoxic state of the skin by competing for oxygen, thereby controlling CD8 T cell-effector function. To test this, we depleted mice of neutrophils for the first two weeks of infection using anti-Ly6G (clone 1A8) (Fig. 5f) at the early stage of disease development and assessed granzyme B expression in CD8 T cells. We found that neutrophil depletion decreased the frequency and the mean fluorescence intensity (MFI) of granzyme B expressing CD8 T cells compared to isotype control-treated mice (Fig. 5g), without changing parasite numbers or lesion size (data not shown). These data suggest that neutrophils consume oxygen in lesions, altering the phenotype of CD8 T cells.

In this regard, activated neutrophils assemble NADPH oxidase to produce reactive oxygen species (ROS), which requires oxygen (75). Thus, we tested whether oxygen consumption to generate ROS resulted in a hypoxic state that impacted CD8 T cell function. For this purpose, we compared the ability of mice sufficient (WT) or deficient in the gp91^phox^ subunit of the NADPH oxidase (Cybb^-/-^) to express granzyme B in lesional CD8 T cells upon infection with *Leishmania*. We found significantly less granzyme B expressing CD8 T cells in Cybb^-/-^ mice than in WT (Fig. 5h), forging a link between NADPH oxidase-dependent ROS production and the development of pathogenic CD8 T cells in cutaneous leishmaniasis lesions.

### The magnitude of hypoxia correlates with the presence of neutrophils in human cutaneous leishmaniasis lesions

To determine whether hypoxia is a feature of human disease, we compared transcriptional signatures in blood and skin from healthy subjects (HS) and cutaneous leishmaniasis (CL) patients using previously published datasets (51, 76). To estimate levels of hypoxia-gene expression in clinical samples, we performed single-sample GSEA using the Harris et al. hypoxia gene signature (58), and referred to this variable as the “hypoxia score” (Supp. Table 3). While there was no difference in the hypoxia scores between blood from cutaneous leishmaniasis patients and healthy subjects (Fig. 6a and Supp. Table 4), the cutaneous leishmaniasis lesions are significantly more hypoxic than healthy skin (Fig. 6b, Data File S3).

**Fig 6.**
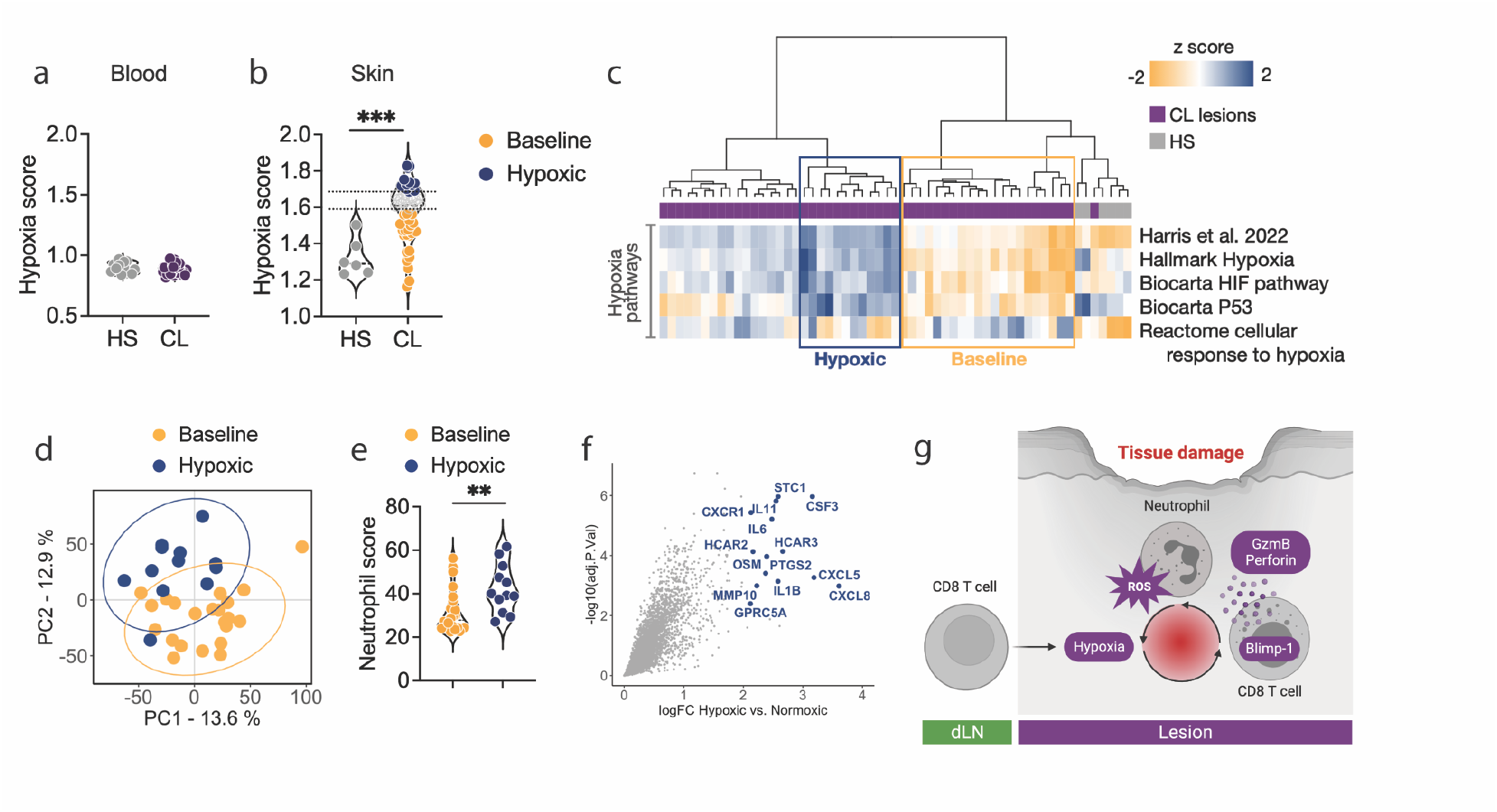
The degree of hypoxia-related gene expression is associated with the presence of neutrophils in patients. (a) Hypoxia score calculated in the blood from healthy subjects (n=14) and patients (n=50); (b) human intact skin (n=6) and cutaneous leishmaniasis lesions from patients (n=51) based on the Harris et al. hypoxia-gene signature. (c) Unsupervised hierarchical clustering classified human skin samples in ‘baseline’ and ‘hypoxic’ lesions according to their hypoxia-gene phenotypes. (d) Principal Component Analysis (PCA) of baseline and hypoxic cutaneous leishmaniasis lesions. (e) Neutrophil abundances estimated by Microenvironment Cell Populations (MCP) counter. Wilcoxon was used for statistical analysis. (f) Scatter plot showing differential gene expression analysis between hypoxic vs. baseline lesions (FDR>0.01 and FC=1.5). In blue are the top DGEs from this analysis. (g) Model summarizing our findings. **P ≤ 0.01, and ***P ≤ 0.001. HS = Healthy subjects. CL = cutaneous leishmaniasis. FDR = false discovery rates. FC = fold change.

Importantly, we observed that the degree of the hypoxia-related gene expression in patients’ lesions was variable. To understand the biological implication of this variability, we performed unsupervised hierarchical clustering (HC) to classify lesions and healthy skin samples according to their hypoxic gene expression accessed by Harris et al., Hallmark, Biocarta and Reactome sources (Fig. 6c). HC analysis revealed two groups of lesion samples: (1) ‘baseline’ lesions (n = 22) with a hypoxia phenotype comparable to control skin samples, and (2) ‘hypoxic’ lesions (n = 12) with a high hypoxic phenotype relative to the entire cohort (Fig. 6b and Fig 6c). The remaining lesions had an intermediate phenotype. PCA revealed that hypoxic lesions are robustly segregated from baseline counterparts (Permanova, Pr(>F)=0.001) (Fig. 6d). Gene Ontology performed for genes overexpressed in hypoxic compared to baseline lesions (FDR>0.01 and FC=1.5 thresholds) revealed statistically significant enrichment for neutrophil chemotaxis and signaling (Supp. Table 5 and Supp. Table 6). Indeed, estimating cell abundances from this unstructured bulk RNA-seq data revealed an increase in neutrophil counts in hypoxic compared to baseline lesions (Fig 6e). The top DEGs between hypoxic compared to baseline lesions included genes associated with neutrophil chemotaxis and survival *CXCL5*, *CXCL8*, *CXCR1*, *CSF3,* and genes encoding for pro-inflammatory cytokines such as *IL1B*, *IL6,* and *OSM* (Fig. 6f and Supp. Table 5). Therefore, our findings in mice are in agreement with the analysis of patient’s samples, implicating the presence of neutrophils to the degree of hypoxia in lesions.

## DISCUSSION

Cytolytic CD8 T cells cause severe inflammation in various settings, such as alopecia, malaria, Steven-Johnson syndrome, toxic epidermal necrolysis, cutaneous leishmaniasis, and many others (14, 15, 21, 24, 31–34, 77). However, the factors that promote cytolysis by CD8 T cells in tissue microenvironments are still poorly understood. Here, we show that neutrophils recruited to the skin upon *Leishmania* infection consume oxygen to produce ROS, generating chronic inflammatory hypoxia. The resultant competition for oxygen changes the aerobic microenvironment and promotes the expression of Blimp-1 in recruited CD8 T cells, which results in the expression of the cytolytic molecule granzyme B. Subsequent cell death within the infected area results in tissue damage and chronic neutrophil recruitment (Fig. 6g). Because there are numerous skin diseases associated with cytotoxicity, our findings provide evidence that local inhibition of this pathway should be considered as a strategy to ameliorate these diseases.

In this regard, preferential expression of cytolytic genes by CD8 T cells in peripheral tissues but not in dLNs is observed in numerous pathogenic conditions. For example, in models of graft-versus-host disease and graft-versus-leukemia, granzyme B is absent in dLNs but present in the skin (9, 11). Additionally, memory CD8 T cells acquire cytolytic activity upon entry into non-lymphoid tissues in vesicular stomatitis virus infections (10). The site-specific function of CD8 T cells is not limited to granzyme B expression. For example, in mice chronically infected with *Trypanosoma cruzi*, CD8 T cells in the spleen produced IFN-γ, while CD8 T cells within the muscle did not (78). Similarly, we found previously that in cutaneous leishmaniasis, CD8 T cells in dLNs produced IFN-γ while CD8 T cells in the skin lacked IFN-γ production (67). Since hypoxic dendritic cells produce less IL-12 (46, 48), this could explain the decrease in CD8 T cell IFN-γ production in lesions (67) and suggests that hypoxia may not only promote pathogenic responses but also prevent protective ones. Conversely, in diseases where cytotoxicity is protective and IFN-γ is pathogenic, the impact of hypoxia would be expected to have the opposite effect. Our confocal microscopy data suggest that hypoxia impacts many other cells beyond CD8 T cells; therefore, it is likely that hypoxia has a much greater role in tissue immunity to *Leishmania*.

There is substantial clinical and experimental evidence that hypoxia contributes to tumor progression, metastasis, and high mortality (79). While low oxygen levels support myeloid suppressor cell accumulation (80) and tumor cell metastasis (81), hypoxia-inducible factors (HIF) provide CD8 T cells with greater anti-tumor potential (82–84). HIF increases CD8 T cell effector function during chronic viral infections and reduces viral titers but leads to higher mortality (85). These findings agree with the observed increases in perforin expression and killing ability of CD8 T cells cultured in hypoxia (84, 86, 87). Similar results were observed in lupus skin disease, where HIF drove granzyme B expression but, in this case, was independent of hypoxia (88). Hypoxia also alters CD8 T cell metabolism (86) and induces tissue residency (82, 89) and exhaustion (90–92) programs. We found a strong correlation between the hypoxic state of CD8 T cells and Blimp-1 expression. Consistent with our findings, hypoxia also triggered Blimp-1 expression in pancreatic tumor cells (81). Since Blimp-1 can instruct tissue residency and exhaustion (62, 93–95), we speculate that hypoxia-driven Blimp-1 expression may also initiate these CD8 T cell programs within cutaneous leishmaniasis lesions, which requires further investigation.

Another immune cell type implicated in the hypoxia-cytotoxicity axis revealed in our study is neutrophils, which rapidly migrate to infection sites and, in many infections, eliminate invading microbes. However, it is well-recognized that the impact of neutrophils extends beyond pathogen killing, as they also shape the function of other cells within tissues by producing cytokines and chemokines, interacting with dendritic cells and T cells, and consequently affecting disease outcomes. For example, neutrophils producing IL-10 (96) or expressing PD-L1 (97) suppress CD4 T cells. Additionally, neutrophils suppress effector responses by tumor- specific T cells in lymph nodes (98) and within tumors (99). In chronic viral infections, suppressive neutrophil subsets reduce the antiviral capabilities of CD8 T cells (100). In contrast, neutrophils facilitate dendritic cell migration to dLNs in contact hypersensitivity and prime allergen-specific CD8 T cells (101). In cutaneous leishmaniasis, neutrophils recruited to lesions can have paradoxical functions depending on the parasite species and mouse model. For example, neutrophils kill *Leishmania* directly by extracellular trap formation or cooperate with macrophages to activate microbicidal activities (102, 103). In contrast, neutrophils reduce costimulatory molecule expression in dendritic cells and silently deliver parasites to macrophages (69, 104, 105). Despite these varied effects, it is clear that chronic recruitment of neutrophils is associated with tissue destruction in cutaneous leishmaniasis (3, 72, 74, 106). Here, we uncovered a new role for neutrophils in cutaneous leishmaniasis lesions by showing that their consumption of oxygen promotes a hypoxic environment, resulting in a phenotypic switch in CD8 T cells.

Cytotoxic CD8 T cells promote increased inflammation by inducing excessive cell death in lesions and enhancing chronic neutrophil recruitment to the skin (3, 4). In what may be related findings, neutrophil depletion decreases the hypoxic state of herpes stromal keratitis lesions and ameliorates disease (107). In a mouse model of colitis, the respiratory burst by Gr1^pos^ cells - encompassing neutrophils and inflammatory monocytes - induced hypoxia in intestinal epithelial cells, which was critical for disease resolution (108). One limitation of our work is that it does not address the impact of other innate cells and their ability to deplete lesions of oxygen, which could also impact CD8 T cell function. However, our work strongly supports oxygen consumption and neutrophil recruitment as modulators of CD8 T cell granzyme B expression, which fosters the chronic presence of neutrophils in lesions. While the negative impact of this cascade of events is clear in cutaneous leishmaniasis, it could be beneficial in other contexts.

To be successful, host-directed therapies for infectious diseases should not augment immune responses that damage tissues and must not interfere with responses that control the pathogen (2). Therefore, it is critical to define immune pathways that are pathogenic rather than protective. In cutaneous leishmaniasis, cytotoxicity is an ideal therapeutic target since it associates with treatment failure in patients who receive anti-parasitic drugs (7) and does not control the parasite (3). Patients affected by cutaneous leishmaniasis are frequently treated with anti-parasitic drugs that require intravenous or intramuscular injections every day over the course of 21 days. Because failure rates are high, many patients must go through two and sometimes three rounds of treatment. Our data provide evidence that local inhibition of the cytolytic program can circumvent the use of systemic drugs, which generate considerable side effects, and topical targets should be considered as a new therapeutic strategy for cutaneous leishmaniasis.

## METHODS

### Mice

C57BL/6 CD45.2 and CD45.1 mice (6 weeks old) were purchased from Charles River, and *Prdm1* YFP (here called Blimp-1 reporter), ODD^cre^/ERT2 (63), Gzmb^cre^, Blimp-1^flox/flox^, and RAG^-/-^ (B6.12957-RAG1^tm1Mom^) were purchased from The Jackson Laboratory and crossed in our mouse facility. Cybb^-/-^ mice (109) were obtained from Dr. Juhi Bagaitkar at Nationwide Children’s Hospital, Columbus, USA. Experiments assessing Blimp-1 expression with Blimp-1 YFP reporter were reproduced using Blimp-1 GFP reporter mice (a kind gift from Stephen Nutt, WEHI, Melbourne, Australia). Gzmb^cre^ mice have the human *GZMB* promoter directing Cre recombinase expression to activated T cells; therefore, it does not directly impact mouse granzyme B expression. Gzmb^cre^ and Blimp-1^flox/flox^ were crossed to generate the deletion of Blimp-1 in activated T cells, and here we called purified CD8 T cells Blimp-1^cKO^. ODD^cre^/ERT2 and Blimp-1^flox/flox^ were crossed to generate mice in which Blimp-1 deletion was restricted to hypoxic cells in tamoxifen-treated mice. All mice were maintained in a specific pathogen-free environment at The Ohio State University or The University of Pennsylvania Animal Care Facilities.

### Parasites

*L. major* (strain WHO/MHOM/IL/80/Friedlin) and *L. braziliensis* (MHOM/BR/01/BA788) were grown in Schneider’s insect medium (GIBCO) supplemented with 20% heat-inactivated FBS (Sigma, St. Louis, MO, USA) and 2 mM glutamine (Thermo-fisher, Waltham, MA, USA). Metacyclic-enriched promastigotes were used for infection (110). Mice were infected with either 10^6^ *L. major* or 10^5^ *L. braziliensis* intradermally in the ear, and the lesion progression was monitored weekly by measuring the diameter of the ear with a digital caliper.

### Cell purification and adoptive transfer

Intralesional cell transfer mouse model: CD45.2 and CD45.1 mice were infected with *L. major*. CD45.2 mice were euthanized three weeks post-infection, and CD8 T cells from dLNs were purified using a magnetic bead separation kit (Miltenyi Biotec, Germany). 10^6^ CD8 T cells were injected into the lesions of CD45.1 mice. Two days post-injection, recipient mice were euthanized, and granzyme B expression in CD8 T cells in the ear and dLN were analyzed by flow cytometry directly ex vivo. RAG^-/-^ mouse model: splenocytes from WT or Blimp-1^cKO^ mice were collected, red blood cells lysed with ACK lysing buffer (LONZA, Switzerland) and CD8 T cells were purified using a magnetic bead separation kit (Miltenyi Biotec, Germany). Three million CD8 T cells were transferred intravenously into RAG^-/-^ mice subsequently infected with *L. braziliensis*. Mice reconstituted with CD8 T cells received four injections of 250 μg of anti- CD4 (clone GK1.5, BioXCell, Lebanon, NH, USA) within the first two weeks.

### In vivo treatment

FTY720 (Cayman Chemical, Ann Arbor, Michigan, USA) was administered intraperitoneally (3 mg/kg) daily 12 days post-infection and for ten days before euthanasia. To induce Cre expression in the ODD^cre^/ERT2 mice, tamoxifen (Sigma, St. Louis, MO, USA) was dissolved in corn oil and administered intraperitoneally (67mg/kg) for seven days following infection. Pimonidazole (Hypoxyprobe Inc, Burlington, Massachusetts, USA) was administered intraperitoneally (9 mg/mL, 200 μL) one hour before euthanasia. For neutrophil depletion, mice received injections every three days of 250 μg of anti-Ly6G (clone 1A8, BioXCell, Lebanon, NH, USA) within the first two weeks.

### Tissue single-cell suspension preparation

Infected ears were harvested, the dorsal and ventral layers of the ear separated, and the ears were incubated in RPMI 1640 (Gibco, Canada) with 250 μg/mL of Liberase TL (Roche Diagnostics, Chicago, IL, USA) for 60-90 mins in a shaker at 37°C and 5% CO2. Ears were dissociated using a cell strainer (40 μm, BD Pharmingen, Franklin Lakes, NJ, USA), and an aliquot of the cell suspension was used for parasite titration. dLNs were homogenized using a cell strainer (40 μm, BD Pharmingen, Franklin Lakes, NJ, USA) to obtain single-cell suspensions.

### Parasite titration

The parasite burden in the ears was quantified as described previously (111). Briefly, the homogenate was serially diluted and incubated at 26**°**C. The number of viable parasites was calculated from the highest dilution at which parasites were observed after seven days.

### In vitro stimulation

dLN single-cell suspensions from C57BL/6 infected mice were cultured in 96 well plates “U” bottom with 5μg/mL of plate-bound anti-CD3 (clone 145-2C11, Invitrogen, Waltham, MA, USA), 0.5 μg/mL soluble anti-CD28 (clone 37.51, Invitrogen, Waltham, MA, USA), 37°C and 5% CO2 for 48 hours in RPMI 1640 containing 100 Units of penicillin and 0.1 mg/mL of Streptomycin (Sigma, St. Louis, MO, USA), 2mM L-Glutamine (Thermo-fisher, Waltham, MA, USA) and 10% FBS (Sigma, St. Louis, MO, USA). Cells were cultured in 1 mM of DMOG (Sigma, St. Louis, MO, USA) dissolved in DMSO or an equivalent volume of DMSO (vehicle) for 48 hours. For mRNA isolation, dLN single-cell suspensions from C57BL/6 infected mice were obtained, and CD8 T cells were purified using a magnetic bead separation kit (Miltenyi Biotec, Germany). CD8 T cells were cultured in 96 well plates “U” bottom with 5μg/mL of plate-bound anti-CD3 (clone 145-2C11, Invitrogen, Waltham, MA, USA), 0.5 μg/mL soluble anti-CD28 (clone 37.51, Invitrogen, Waltham, MA, USA), 20 U/mL recombinant human IL-2 at 37°C and 5% CO2 for 24 hours in RPMI 1640 (Gibco, Canada) containing 100 Units of penicillin and 0.1 mg/mL of Streptomycin (Sigma, St. Louis, MO, USA), 2mM L-Glutamine (Thermo- fisher, Waltham, MA, USA) and 10% FBS (Sigma, St. Louis, MO, USA). Cells were cultured in 1 mM of DMOG (Sigma, St. Louis, MO, USA) dissolved in DMSO or an equivalent volume of DMSO (vehicle).

### Flow cytometric analysis

Before surface and intracellular staining, cell suspensions were stained with a LIVE/DEAD fixable aqua dead cell stain kit (Thermo-fisher, Waltham, MA, USA), according to manufacturer instructions. Analysis was performed using the FlowJo Software (Tree Star, Ashland, OR, USA), and gates were drawn based on fluorescence minus one control. List of antibodies: CD45 (clone 30-F11), GzmB (clone GB11), CD11b (clone M1/70) (all Invitrogen, Waltham, MA, USA). CD44 (clone IM7, BD Biosciences, Franklin Lakes, NJ, USA), CD45.2 (clone 104), CD45.1 (clone A20), CD90 (clone 53-2), CD3 (clone 17A2), CD8β (clone YTS156.7.7), and Ly6G (clone 1A8) (all from BioLegend, San Diego, CA, USA). The stained cells were run on BD FACSymphony A3 or FACSCanto II (BD Biosciences, San Jose, CA, USA).

### Quantification of *Prdm1* mRNA by qPCR

RNA extraction of in vitro stimulated CD8 T cells was performed using nulceospin RNA, Mini kit (Macherery-Magel, Germany) and used to prepare complementary DNA (cDNA) using iScript Reverse Transcription Supermix for RT-qPCR (Bio-Rad, Hercules, CA, USA) on ABS Proflex Thermal cycler (Thermo-fisher, Waltham, MA, USA). qPCR was carried out on a C1000 RT-PCR using iTaq Universal SYBR Green Supermix (all from Bio-Rad, Hercules, CA, USA) and primers targeting *Prdm1* (forward 5’-TTC TCT TGG AAA AAC GTG TGG G-3’ and reverse 5’-GGA GCC GGA GCT AGACTT G-3’). The qPCR results were normalized to β- Actin (forward 5’-CGC TGT ATT CCC CTC CAT CG-3’ and reverse 5’-CCA GTT GGT AAC AAT GCC ATG T-3’). All reactions were carried out in duplicates, and data is represented as fold-change (FC) over the average expression of vehicle-treated cells.

### Histology

Two weeks after infection, mice were euthanized, and infected skin (lesions) and contralateral (naïve skin) ears were collected, immediately immersed in 2% paraformaldehyde (PFA) (Thermo-fisher, Waltham, MA, USA), and incubated at room temperature for one hour. Ears were divided in half and re-immersed in PFA for another hour at room temperature.

Samples were rinsed and immersed in phosphate-buffered saline (PBS, pH 7.4) (Gibco, Canada) for two hours, followed by cryoprotection in sucrose 30% for 48 hours. Cryoprotected samples were preserved in optimal cutting temperature (OCT) compound (Thermo-fisher, Waltham, MA, USA), and serial sections of 15 µm were obtained on a Tissue-Tek Cryo3 Flex cryostat (Sakura Finetek, Torrance, CA). Sections were collected onto Superfrost Plus microscope slides (Thermo-fisher, Waltham, MA, USA), air-dried, and stored at -80^°^C.

### Immunofluorescence, Image acquisition, and Analysis

Slides were dried at room temperature for one hour, rinsed in PBS, and incubated for one hour in blocking solution (4% bovine serum albumin and 0.1% Triton X-100 (both from Thermo-fisher, Waltham, MA, USA) dissolved in PBS). Sections were incubated overnight at 4^°^C with rabbit anti-Pimonidazole antisera (Hypoxyprobe Inc, Burlington, Massachusetts, USA). The slides were washed with PBS and incubated for 4 hours at room temperature with rat anti-mouse Ly6G (cone 1A8, BioLegend, San Diego, CA, USA). Slides were washed with PBS and incubated separately with Alexa Fluor 568 goat anti-rabbit (Invitrogen, Waltham, MA, USA) and Alexa Fluor 488 goat anti-rat (Invitrogen, Waltham, MA, USA). Sections were covered in each secondary antibody for two hours at room temperature. Finally, the sections were counter-stained with the nuclear probe 4’,6-diamino-2-phenylindole (DAPI, Biotechne-Tocris, UK). Slides were covered with Prolong™ Gold Antifade Mountant (Invitrogen, Waltham, MA, USA) and kept protected from the light at room temperature for a minimum of 24 hours to allow the mounting media to dry before imaging. Representative images were acquired with 10x and 20x objectives in an Olympus FV3000 confocal microscopy (Olympus Corporation, Japan). The color of the different structures was digitally inverted to better match the presented data in this manuscript.

Using ImageJ software (version 1.53, National Institute of Health), 27 regions of interest (ROIs) measuring 600x40 µm were created. From the low signal edge to the other, the pixel intensity values of each ROI were obtained.

### Transcriptional profiling of purified mouse CD8 T cells

C57BL/6 mice were infected with *L. major,* and five weeks post-infection, mice were euthanized; cells from the infected ear and dLN cells were purified and stained for flow cytometric analysis. Antigen-experienced CD8 T cells were purified using FACSAria cell sorter (gating strategy: live, singlets, CD90^+^, CD8^+^CD44^high^), and RNA was extracted using the RNeasy Plus Mini Kit (QIAGEN, Germany) according to the manufacturer’s instructions and used to prepare Poly(A)+-enriched cDNA libraries Illumina TruSeq Stranded mRNA library prep workflow. Ribo-Zero Gold rRNA depletion (Illumina, San Diego, CA, USA) removed ribosomal content. Quality assessment and quantification of RNA preparations and libraries were performed using an Agilent 4200 TapeStation and Qubit 3, respectively. Samples were sequenced on an Illumina NextSeq 500 to produce 75–base pair single-end reads. The raw reads were mapped to the mouse reference transcriptome (Ensembl; Mus musculus version 108) using Kallisto version 0.46.0, and MultiQC v1.9 was used to check the quality of the alignment.

All subsequent analyses were conducted using the statistical computing environment R version 4.1.0, RStudio version 1.4.1717, and Bioconductor version 3.13. Briefly, transcript quantification data were summarized to genes using the BiomaRt and tximport package and normalized using the trimmed mean of M values (TMM) method in edgeR. Genes with <1 CPM in at least three samples were filtered out. Normalized filtered data were variance-stabilized using the voom function in limma, and Differential Gene Expression analysis was performed with linear modeling using limma after correcting for multiple testing using Benjamini-Hochberg FDR correction. The list of 7 genes encoding for transcription factors associated with effector like T cell functions was gathered from (55–57). Hypoxic enrichment per RNA-seq sample was calculate with single-sample GSEA using the GSVA R package. The 5 Hypoxia-related signatures used to estimate those levels were downloaded from: Harris et al. 2022 (58), Hallmark systematic name (sn) #M5891, Biocarta sn #M13324, Biocarta sn #M14863 and Reactome sn #M641.

### Hypoxia-related transcriptional analyses on lesion biopsies from cutaneous leishmaniasis patients

Data in Fig. 6 are derived from published transcriptional profiling (51, 76). The analysis carried out for this current study was performed from the filtered, normalized gene expression matrix available for download as a text file on NCBI GEO accession #GSE162760 and #GSE214397. All subsequent analyses were conducted similarly to the purified mouse CD8 T cells RNA-seq dataset. Hypoxia scores for the human RNA-seq dataset were calculated using the Harris et al. gene signature (58). Unsupervised hierarchical clustering classifying lesion samples according to their “hypoxic scores” was performed with maximum distances and Ward’s D2 minimum variance. Permanova statistical testing was performed with the vegan R package. Gene Set Enrichment Analysis (GSEA) was performed using the Broad Institute software version 4.3.2. Gene Ontology (GO) analyses were carried out using DAVID Bioinformatics Resources (2021 Update) from NIAID/NIH and biological process terms, clustering genesets by annotation similarity. MCP-counter and immundeconv R packages were combined to estimate neutrophil abundances from the unstructured RNA-seq dataset. Data visualization was mostly performed in GraphPad Prism version 10 and in R programming language using tidyverse and gplots tools.

### Statistics

Statistical significance was determined using the two-tailed unpaired Student’s *t*-test. The specific transcriptional analysis section describes detailed statistical analysis for RNA sequencing. Differences were considered significant when *p* ≤ 0.05 (*), *p* ≤ 0.01 (**), *p* ≤ 0.001 (***), or *p* ≤ 0.0001 (****).

## Study Approval

This study was carried out per the recommendations in the Guide for the Care and Use of Laboratory Animals of the National Institutes of Health. The Institutional Animal Care and Use Committee, The Ohio State University, and The University of Pennsylvania approved the protocol.

## Data availability

Raw sequence data from purified mouse CD8 T cells are available on the NCBI GEO BioProject PRJNA794119. The complete RNA-seq data analysis, R code, file inputs, and outputs used to perform the transcriptional analysis presented in this current manuscript are available on the GitHub repository “Hypoxia_FON” (https://github.com/camilafarias112/Hypoxia_FON).

## Author contributions

EAF, CFA, RAS, EDSH, KM, LAS, WZ, CIO, GDW, PLC, and FON designed research studies; EAF, CFA, RAS, EDSH, KM, LAS, WZ, FON, and PLC conducted experiments; EAF, CFA, KAM, WZ, and FON analyzed and interpreted the data; EAF, CFA, and FON wrote the manuscript. FON supervised the study.

## Acknowledgments

This work was supported by National Institutes of Health grant R01AI162711 (FON), National Institute of Health training program T32 AI165391 “Interdisciplinary Program in Microbe-Host Biology” (EAF), National Institute of Health training program T32 AI118684 (WZ), Host-Microbial Analytic and Repository Core in the Center for Molecular Studies in Digestive and Liver Diseases (P30DK050306) (GDW). The authors would like to thank Dr. Juhi Bagaitkar for providing Cybb^-/-^ mice.

## Supplemental Material

**Supp. Fig. 1.**
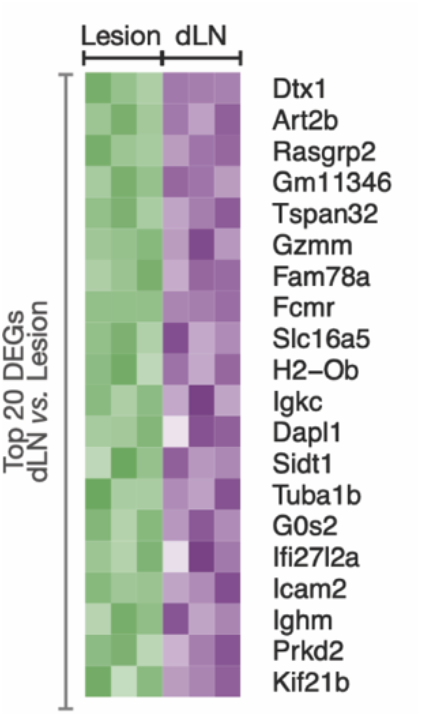
Genes overexpressed in CD8 T cells purified from dLNs compared to lesions. C57BL/6 mice were infected with *L. major*, and at the peak of the lesion, antigen-experienced (CD44^high^) CD8 T cells were purified by flow cytometry cell sorting and used for RNA-seq. Heat map of top 20 differentially expressed genes overexpressed in CD8 T cells purified from dLN compared to lesions.

**Supp. Fig. 2.**
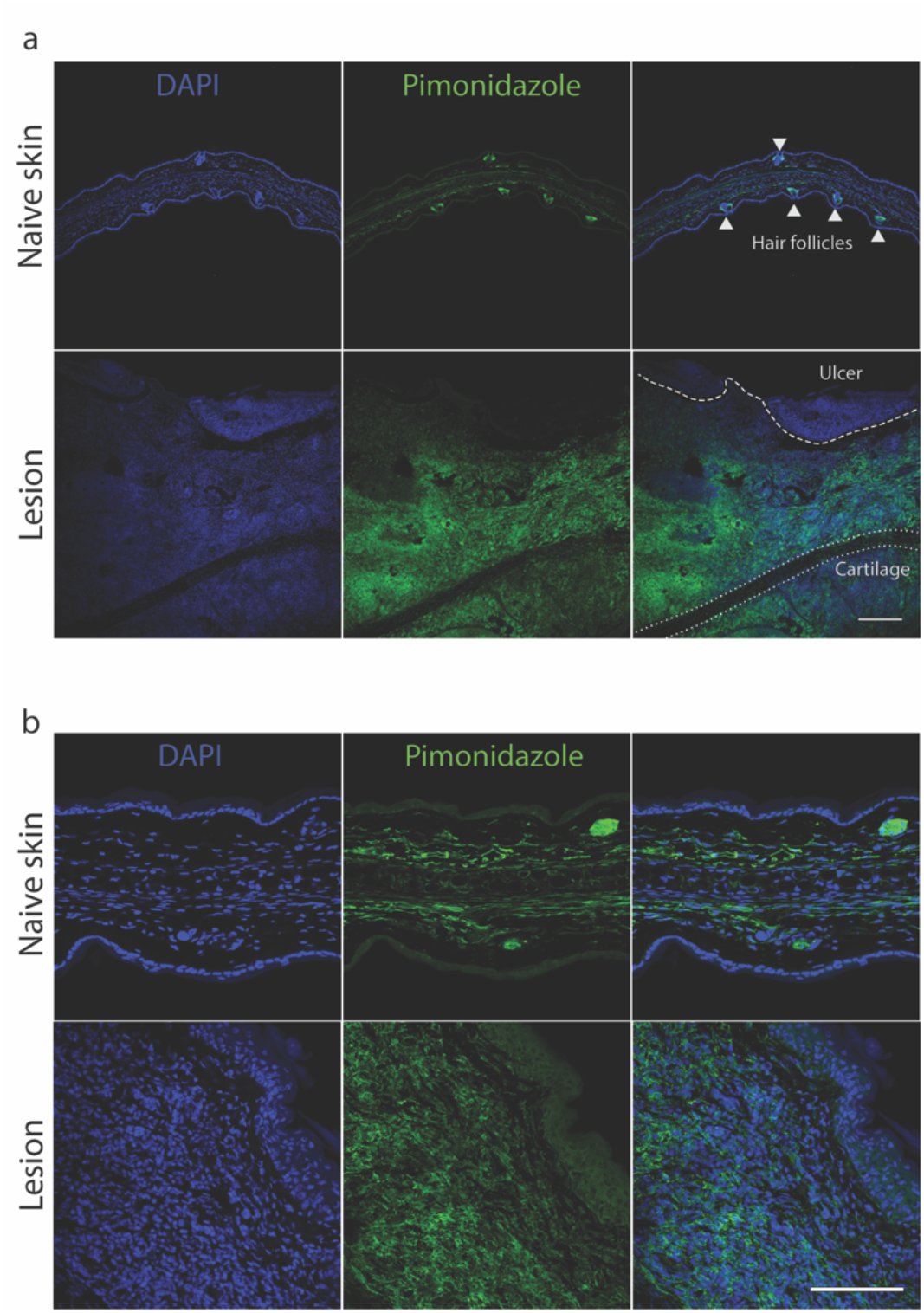
Pimonidazole expression in *Leishmania*-infected skin. C57BL/6 mice infected with *L. major* received pimonidazole one hour before euthanasia at two weeks. (a) Individual images from confocal microscopy (Fig. 2a). Nuclear (DAPI) staining (blue) and pimonidazole (green) in contralateral ears (naïve skin) and infected ears (lesions) for two weeks. Scale bar = 200 μm. (b) Individual images from confocal microscopy (Fig. 2b). Nuclear (DAPI) staining (blue) and pimonidazole (green) in contralateral ears (naïve skin) and infected ears (lesions) for two weeks. Scale bar = 100 μm.

**Supp. Fig. 3.**
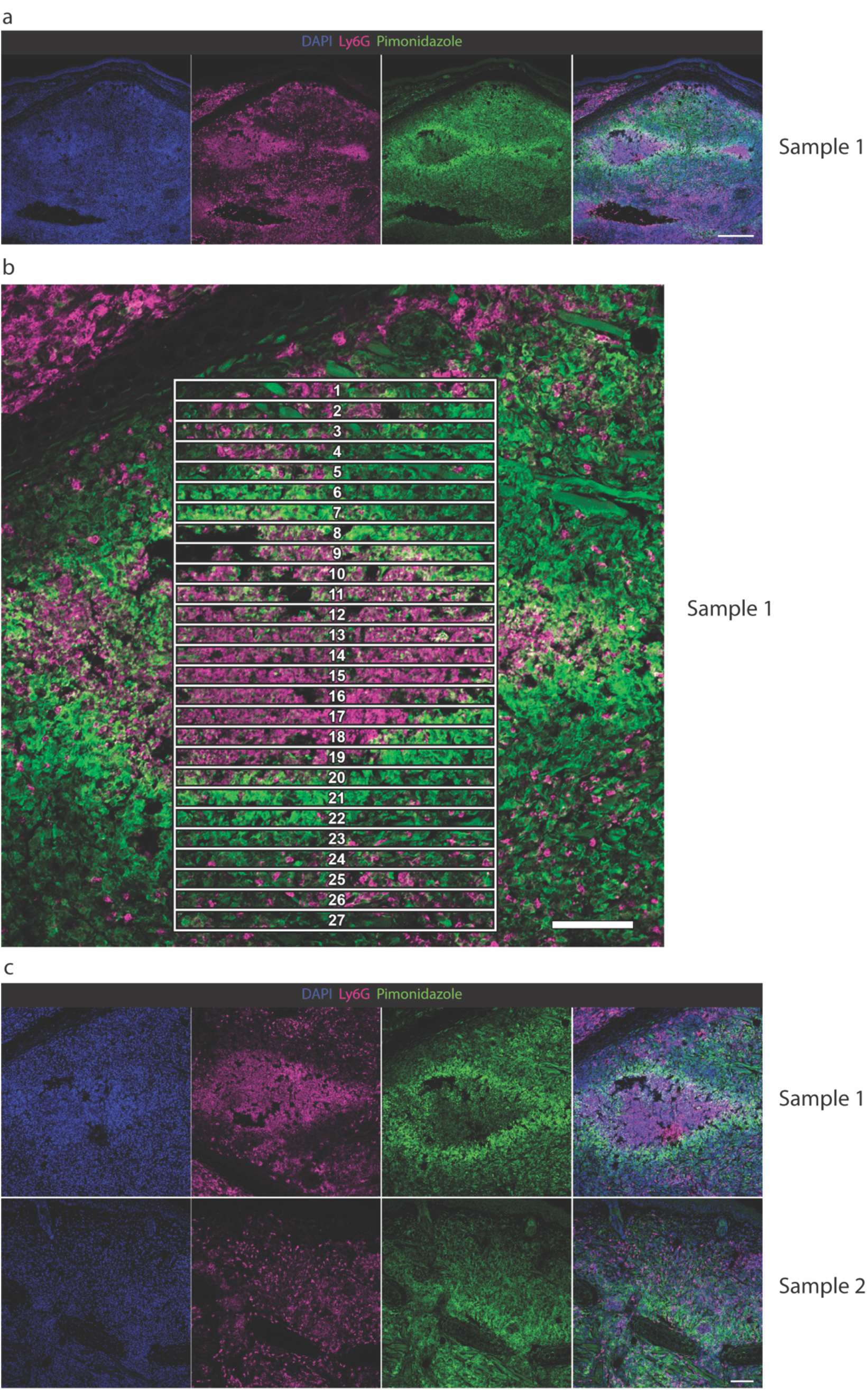
Neutrophil and pimonidazole expression in *Leishmania*-infected skin. C57BL/6 mice infected with *L. major* received pimonidazole one hour before euthanasia at two weeks. (a) Individual images from confocal microscopy (Fig. 5c). Nuclear (DAPI) staining (blue), Ly6G (pink), and pimonidazole (green) in the skin infected for two weeks. Scale bar = 200 μm. (b) Confocal microscopy image was divided into 27 regions to measure the pixel intensity of Pimonidazole (green) and Ly6G (pink) shown in Fig. 5d. Scale bar = 100 μm. (c) Individual images from confocal microscopy (Fig. 5e). Nuclear (DAPI) staining (blue), Ly6G (pink), and pimonidazole (green) in the skin infected for two weeks. Scale bar = 100 μm.

